# Survey-Based Analysis of a Science of Science Communication Scientific Interest Group: Member Feedback and Perspectives on Science Communication

**DOI:** 10.1101/2024.06.25.600615

**Authors:** Anna J. Hilliard, Nicola Sugden, Kristin M. Bass, Chris Gunter

## Abstract

Coordinated attempts to promote systematic approaches to the design and evaluation of science communication efforts have generally lagged behind the proliferation and diversification of those efforts. To address this, we founded the US National Institutes of Health (NIH) Science of Science Communication Scientific Interest Group (SciOSciComm-SIG) and undertook a mixed-methods survey-based evaluation of the group one year after its founding. Respondents indicated ongoing interest and some participation in public-facing science communication while identifying specific barriers, and praised the role of the SIG in expanding access to information about evidence-based practices.

## Introduction

Staff and students at all types of research institutions—universities, government, nonprofits, and private industry—are no strangers to the mounting pressure for scientists to communicate their work to the general public. In response, science communication activities have proliferated across research environments, providing opportunities for interdisciplinary collaboration and greater engagement with non-scientists [McKee et. al, 2022; Naughton et. al, 2024]. Recent science communication efforts continue to make use of traditional channels like television appearances and print media alongside more modern online platforms and experimental approaches [Entradas et al., 2020]. An expansion in the development and availability of specialized trainings has accompanied the rise in science communication practice, as have limited analyses of the efficacy of such trainings, which have raised concerns over the consistency, oversight, and impact of science communication training methods [Baram-Tsabari & Lewenstein, 2017; Rubega et. al, 2021].

Broadly, the progress in science communication efforts and training opportunities has not been matched by efforts made in the systematic design and evaluation of these ventures. The scarcity of so-called “evidence-based” science communication has been attributed to a variety of multifactorial phenomena. Jensen & Gerber [2020] propose a leveled hierarchal framework describing a disconnect between research and practice in the field of science communication, ranging from challenges in meaningfully determining the relevance or applicability of findings in science communication research to ensuring that these research findings are accessible and methodologically robust. Evaluation efforts that exist in spite of research-practice disconnect are not always publicly available, and those that are have been found to often take the role of telling “success stories” that represent the end of a particular science communication venture rather than a foundation of evidence for other communicators to utilize and build upon [Jensen, 2015; Ziegler et al., 2021]. Additionally, many of those communicating science are not themselves scholars in communication, leading to a disconnect in expectations between those trying to communicate and those evaluating that work [Yuan, Besley, & Dudo, 2019]. Regardless of discipline, field, or setting, there is a clear insufficiency in attempts to evaluate science communication efforts.

In the United States (US), one of the main venues for federal-level engagement in science communication of biomedical research is the National Institutes of Health (NIH), who share content through many avenues including online channels, briefings for other areas of government, press interactions, working with advocacy groups, and traditional academic outputs. Like any large and complex institution, the professional environment of the NIH presents its own unique challenges for science communication, and these serve quite different objectives. For context, the NIH is made up of a collection of institutes, centers, and offices (ICOs), most serving as home to a number of research groups (intramural research), while each ICO also sends the bulk of its funds to investigators in other institutions (extramural research); naturally, the ICOs often want to communicate about all of it. The NIH Office of the Director may focus science communication efforts on increasing institutional visibility by highlighting specific health and science advancements funded by the NIH or reaching those who control the funding for the institution [“How the NIH Brings Health,” 2018]. In contrast, individual laboratories may be most interested in reaching very small, specific communities to encourage participation in their own research, such as for rare conditions [Peters, 2022]. To facilitate this, NIH has significantly invested in encouraging individual investigators to learn better science communication through workshops, trainings, resources, and other similar training opportunities [“Clear Communication,” 2021].

Differing objectives, of course, require different communication strategies and different metrics to define and judge success. Therefore, it is crucial that staff looking to use the most effective approaches to fulfill any of these communication objectives are appraised of the latest work in science communication research, and have access to information about a range of methods for the design and evaluation of science communication activities.

The Science of Science Communication Interest Group (SciOSciComm-SIG) [“Science of Science Communication,” 2024] was formed in July 2022 to bring together NIH staff and trainees interested in creating and applying systematic approaches to the design and evaluation of science communication efforts. Monthly meetings in the form of online seminars and journal clubs focus on measures of effectiveness and methods to increase the success of science communication efforts. By July 2023, the group had over 500 members on its mailing list, and regular attendance of over 100 members at its monthly meetings. To fulfill the mission of providing wider exposure to scholarship in related areas, our SIG has included guided discussions on research priorities for health misinformation on social media, seminars on the effects of uncertainty in public science communication, the effects of retractions on scientific publishing and metrics, and other important topics [Chou, Gaysynsky, & Cappella, 2020; Gustafson and Rice, 2020; Oransky, 2022]. As we approached the one-year anniversary of the SIG’s founding, we recognized an opportunity to evaluate the work of the SIG thus far as well as the current scope of science communication activities within the NIH. To broaden the scope of insight gained from study participants and minimize the effects of sampling bias, we additionally sought out the perspectives of NIH staff who are not members of the SIG, herein referred to as “non-members.”

We sought to explore 1) what differences in science communication activity frequencies and perspectives on science communication, if any, exist between SIG members and non-members; 2) the current landscape of science communication activities within the NIH and what barriers, if any, are present to science communication; and 3) SIG member perspectives on the past and present activities of the SIG as well as its future directions.

## Methods

### Survey Design and Distribution

We carried out an anonymous, online mixed-methods survey to collect information on communication activity types and frequencies within the last 12 months, opinions on NIH-specific barriers to science communication, preferences for future SIG activities, respondent demographics, and other similar metrics. We distributed two versions of the survey: one version for SIG members that had additional questions regarding respondents’ experiences with the SIG, and a non-member version that omitted member-specific questions. Responses were mandatory for all survey questions to ensure complete data. The survey, with an estimated completion time of 10 minutes, was hosted on Qualtrics and was distributed to participants via email. We provided participants compensation in the form of a digital $10 Amazon gift card. Our research protocol was deemed exempt by the NIH Office of Research Protections.

### Participants and Screening

Participants from the SciOSciComm-SIG, referred to hereafter as “members,” were recruited via emails sent to the SciOSciComm-SIG LISTSERV. Participants from the NIH Immunology and Bioinformatics SIGs, referred to as “non-members,” were similarly recruited via emails sent to their corresponding SIG listservs. Per exempt research protocol regulations, participant consent was acquired via a text-based consent script to be read and agreed to by participants immediately prior to beginning of survey. To be eligible, participants had to be NIH personnel, at least 18 years of age, and able to read English.

### Science Communication Activity Measures

To capture the types and frequencies of science communication activities respondents were involved in, we adapted a pre-existing measure [Entradas et al., 2020] that assessed the frequency of each respondent’s participation in a list of different science communication activities in the 12 months prior to participation in the survey. In our revised measure, science communication activities were separated into four categories, three of which are public-facing (i.e. meant for a general audience) and one of which is academic-facing (i.e. meant for fellow specialists), and each of which consisted of a varying number of sub-categories representing a specific science communication activity (Table 1). The three public-audience science communicating categories included “Events,” which consists of large-scale public science events such as public workshops, “Traditional Media,” which includes media-based activities such as appearances in televised programming, and “Online,” which encompasses a wide variety of online-based science communication activities, including posts on various social media platforms. The sole academic-facing category, “Academic,” is comprised of common specialist-facing communication activities such as presenting a scientific poster at a conference.

**Table 1.**
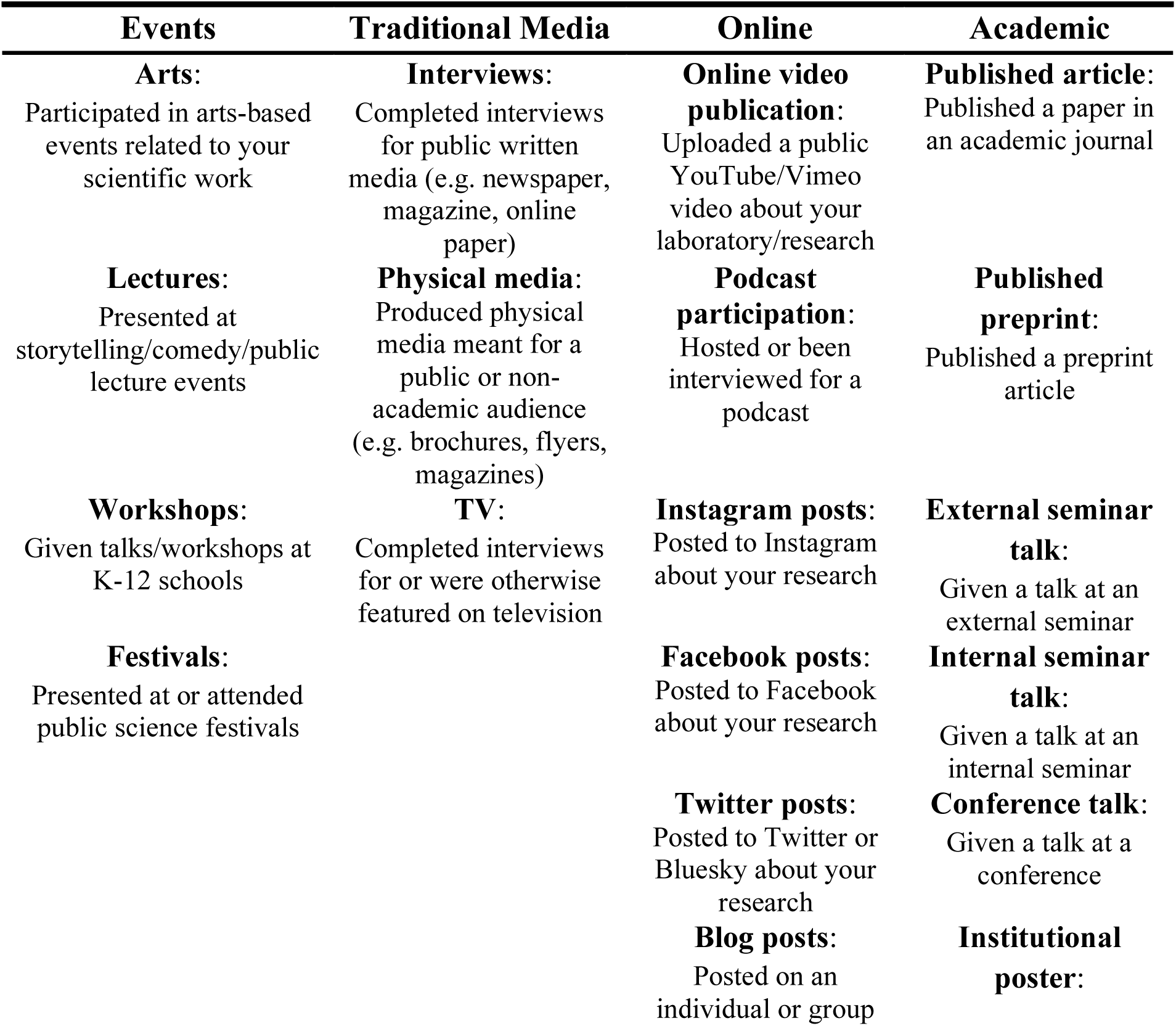
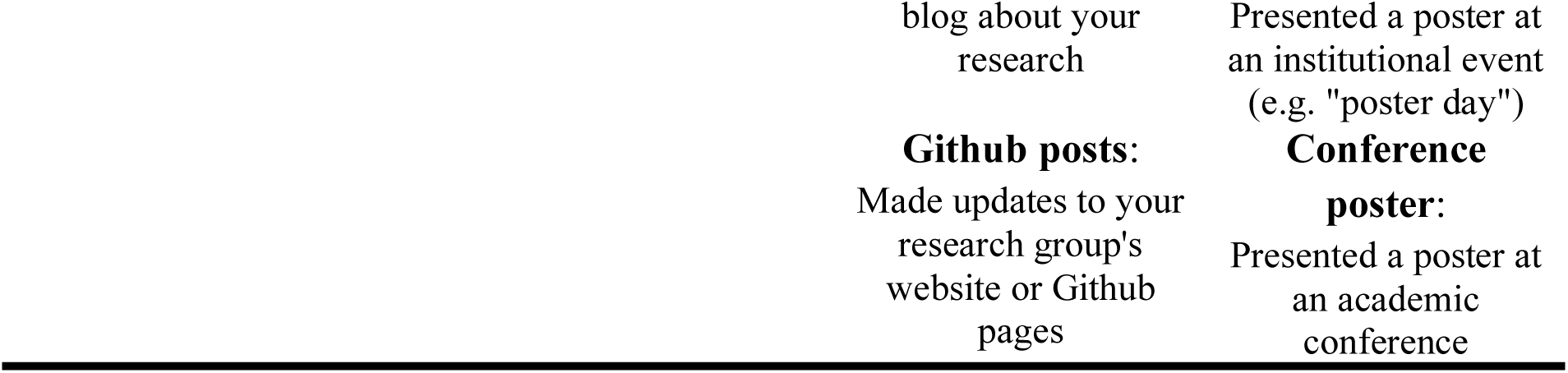
Items included under each of the four science communication frequency subcategory scales. Item descriptions match those provided in the survey.

To account for activities outside of the scope of these pre-defined categories, we additionally asked about the frequency of *any* type of public-facing and academic-facing science communication activities that occurred in the last 12 months. In line with Entradas *et al*, all science communication frequency scales were framed in the context of the respondent’s research group, defined in the survey as, “the laboratory or unit, generally led by one or more principal investigators, that [the respondent is] most closely affiliated with.” We asked respondents who felt that they were affiliated with multiple research groups to base their responses on the group that they would consider to be their primary research unit.

The science communication frequency measures were contextualized using a measure of perceived success, for which participants were asked to rate how effective or successful they felt their science communication efforts to be on a five-point Likert scale, with options ranging from 1 (Not at all successful) to 5 (Very successful).

### Assessment of Perspectives on and Barriers to Science Communication

To gain an understanding of participants’ perspectives on science communication, particularly its goals and ideal success metrics, we asked two open-ended free response questions with no character limit. Firstly, we asked, “In your experience, what are the most effective metrics for measuring the success of science communication activities” Secondly, participants were asked, “What do you think is the most important goal of science communication?”

Additionally, we asked participants about NIH-specific barriers to science communication efforts in two formats: an initial open-ended question with no character limit that asked, “What barriers to science communication, if any, have you faced while at the NIH?” and a second question that was in a multiple-choice format asking participants to identify the biggest barrier to science communication at the NIH. The response options for the multiple-choice question included four predetermined options that the authors predicted would be the most common among respondents, as well as a fifth “Other” option that allowed for free-text response (see Figure 7). To assess the role of the SIG in addressing barriers to science communication, we asked a free-response question reading, “What, if any, barriers to science communication has the SciOSciComm-SIG helped to alleviate or address for you or your research group?”

### Member Feedback on the SIG and Its Impact on Practice

We assessed the impact of the SIG on member work practices by asking SIG member participants, “What knowledge, techniques, approaches, etc. highlighted in SciOSciComm-SIG meetings, if any, have you applied in your own work?” in a free-text open response format. Furthermore, to gather feedback on the present and future structure and activities of the SIG, member respondents were provided a matrix of potential changes made to the group and asked to rate the impact of each change on their likelihood of future SIG meeting attendance on a five-point Likert scale ranging from “Much less likely to attend” to “Much more likely to attend.” Hypothetical changes to the SIG included: increased or decreased meeting frequency, holding more or fewer general body/forum meetings, holding journal club-style meetings, diversifying meeting speaker topics, allowing members to nominate speakers, and polling SIG members about desired speakers or topics.

### Demographics and NIH-Specific Descriptors

We collected basic demographic information from participants, both personal (e.g. gender identity) and occupational; for the latter, several of the metrics were specific to NIH-based positions. For example, we asked about participants’ occupational roles, with the options of “Intramural,” meaning that they are directly affiliated with a scientific laboratory within the NIH; “Extramural,” i.e. involved in providing grant funding to institutions outside of the NIH; and “Administrative,” which encompasses a variety of research-supporting roles. We also asked about respondents’ institute, office, or center (ICO) affiliation, as the NIH is divided fiscally and managerially into ∼28 primary ICOs. Additional demographic information collected included participant research area (if involved in research), which we divided into the pre-existing NIH Intramural Research Program research categories, and training level/career status.

### Statistical Analysis

We analyzed and visualized quantitative data using R/RStudio, Microsoft Excel, and SPSS software. We measured statistical significance using unpaired t-tests for parametric data and Mann-Whitney U-tests for nonparametric data with α = 0.05, adjusted via Bonferroni correction where appropriate.

Thematic analysis of qualitative data was undertaken in Excel by two coders through an inductive methodology. Coder 1 first created a set of code categories for each open-response question based on general themes seen in the responses, and subsequently assigned individual responses to relevant codes. Coder 2 created a revised framework (consolidating, adding, or removing code categories) using Coder 1’s framework as a base, and then coded participant responses appropriately. Disagreements between the two coders were either manually resolved by Coder 1 (in the case of objectively incorrect categorizations, data shifts, etc.) and were ultimately resolved to full agreement through discussion. We represented the frequency of a given code as the percentage of comments attributed to that code out of the total number of comments in response to the relevant prompt, and produced visualizations of coded comments in Excel.

## Results

### Demographics

Respondents from both the SIG member and non-member groups predominantly identified as women (88% and 65%, respectively), and one participant in the member group identified as non-binary (Table 2). One respondent in each group selected “Other” as their gender option and indicated in a free-text response that they preferred not to answer the question. Participant career levels ranged from post-baccalaureate trainees up through principal investigators or equivalent administrative positions. Member group respondents were fairly evenly distributed between the three types of occupational role, whereas most non-member participants (68%) indicated that they work in an intramural role (Figure S1).

**Table 2.** Self-described gender identities of survey respondents.

|  | Woman<br>*n* (%) | Man<br>*n* (%) | Non-binary<br>*n* (%) | Other<br>*n* (%) | Total |
| --- | --- | --- | --- | --- | --- |
| **Member** | 46 (88) | 4 (8) | 1 (2) | 1 (2) | 52 |
| **Non-member** | 26 (65) | 13 (32.5) | 0 (0) | 1 (2.5) | 40 |

Approximately 62% of SIG members and 95% of non-members indicated that they are directly involved with research as a part of their occupational role (Figure 1). Both groups covered a wide variety of NIH, spanning 16 of the 22 NIH intramural research areas and 18 of the 28 ICOs (Figure 1; Figure S2). SIG members demonstrated greater diversity in ICO than non-members, likely due to the comparison SIGs being focused on specific scientific fields (Figure S2).

**Figure 1:**
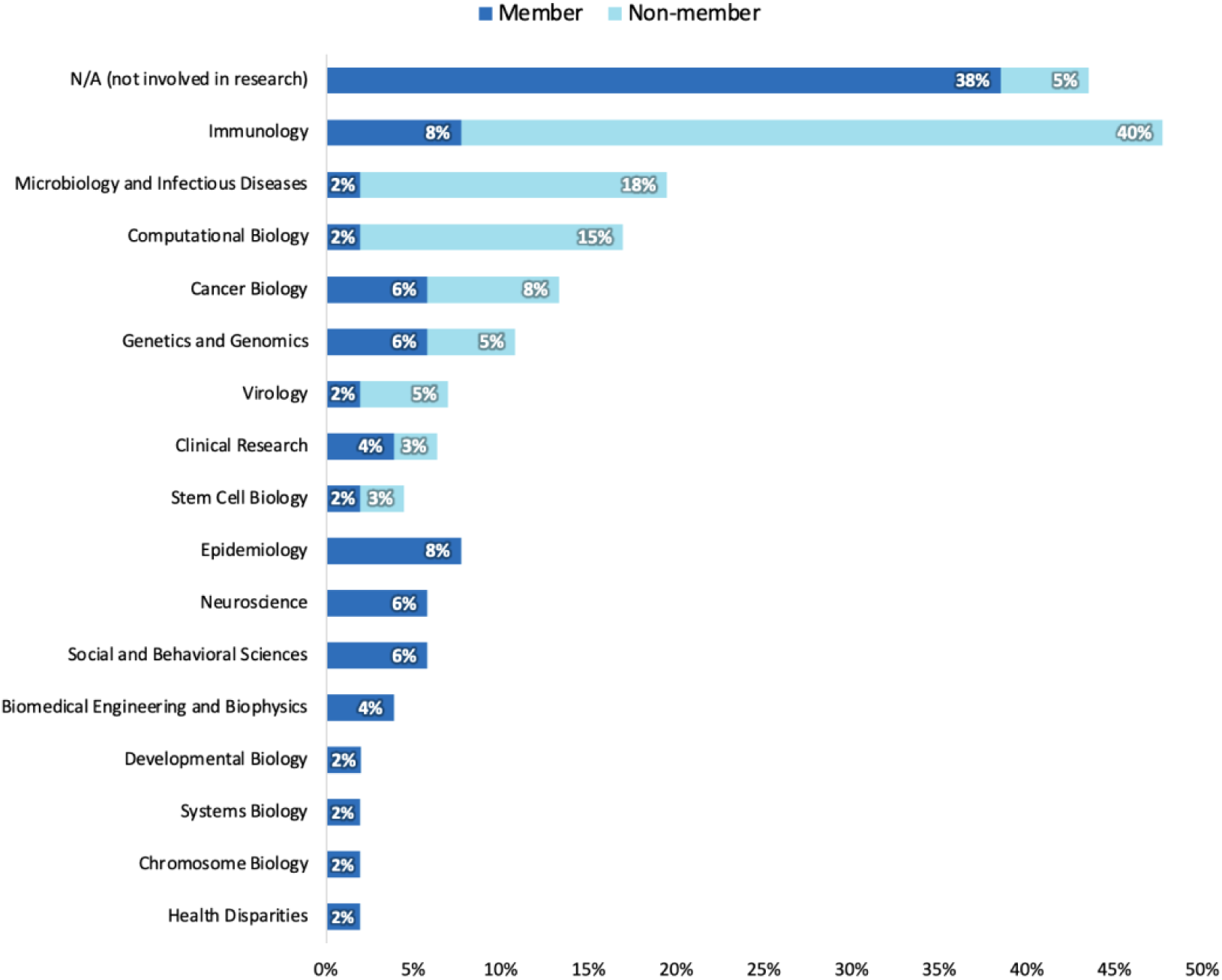
Primary research affiliation for SIG member (*n* = 52) and non-member (*n* = 40) survey respondents.

### Perspectives on Science Communication

To better contextualize the science communication activity data provided by our respondents, we additionally asked both SIG members and non-members to describe what they feel to be the most important goal of science communication activities as well as what they believe to be the most effective metrics for measuring the success of science communication efforts. Inductive thematic analysis revealed a wide variety of goals and metrics: SIG members most commonly mentioned ensuring that their science is clear, comprehensible, and accessible to its target audience as the most important goals of science communication (Figure 2A). SIG members also seemed to favor the implementation of qualitative metrics for the success of science communication efforts to a greater extent than non-members, and were also more likely to prioritize audience engagement as a success metric (Figure 2B).

**Figure 2A:**
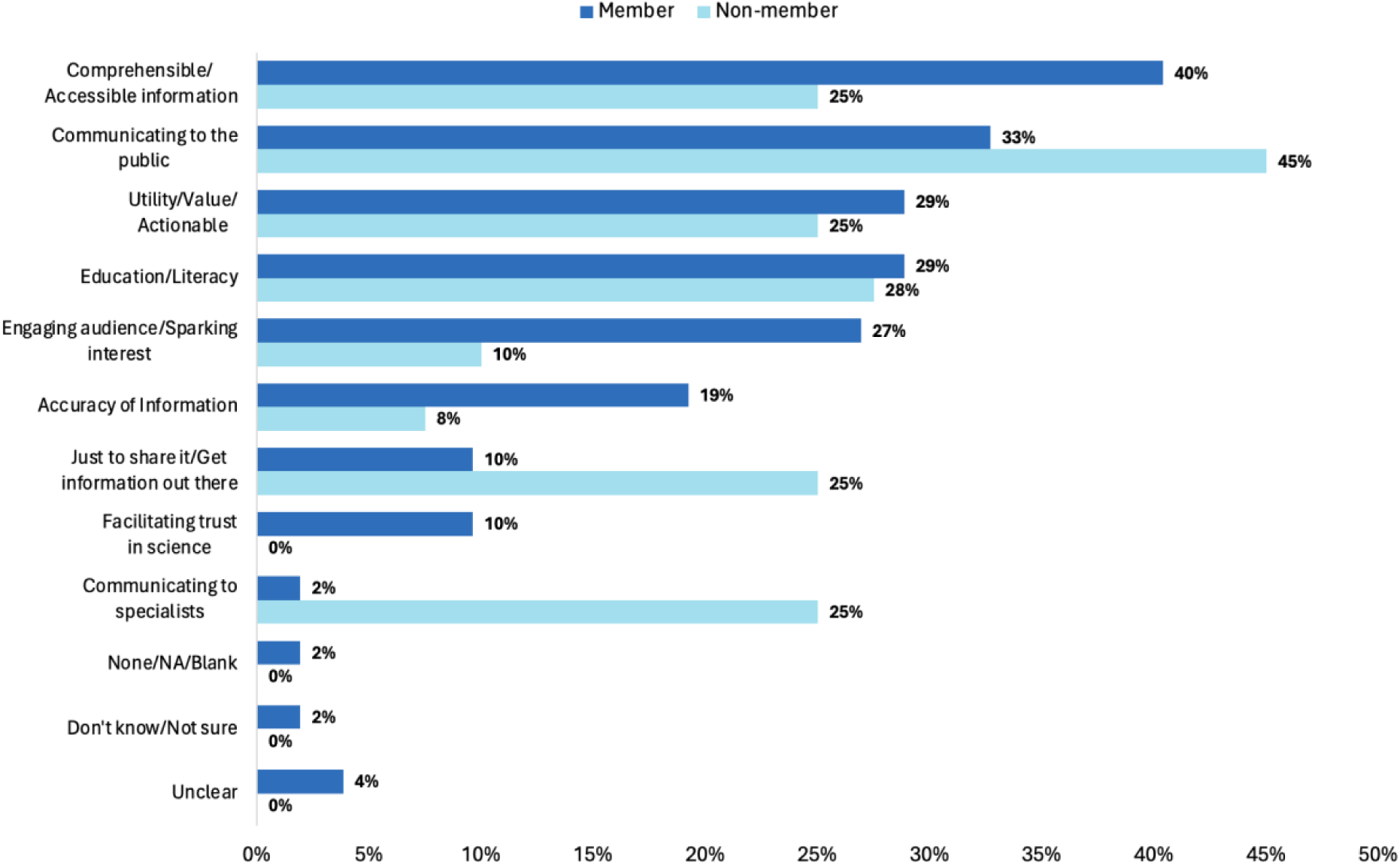
Qualitative coding of SciOSciComm-SIG member (*n* = 52) and non-member (*n* = 40) responses to what they felt to be the most important goal of science communication. Respondents were provided an open response text box with no character limit.

**Figure 2B:**
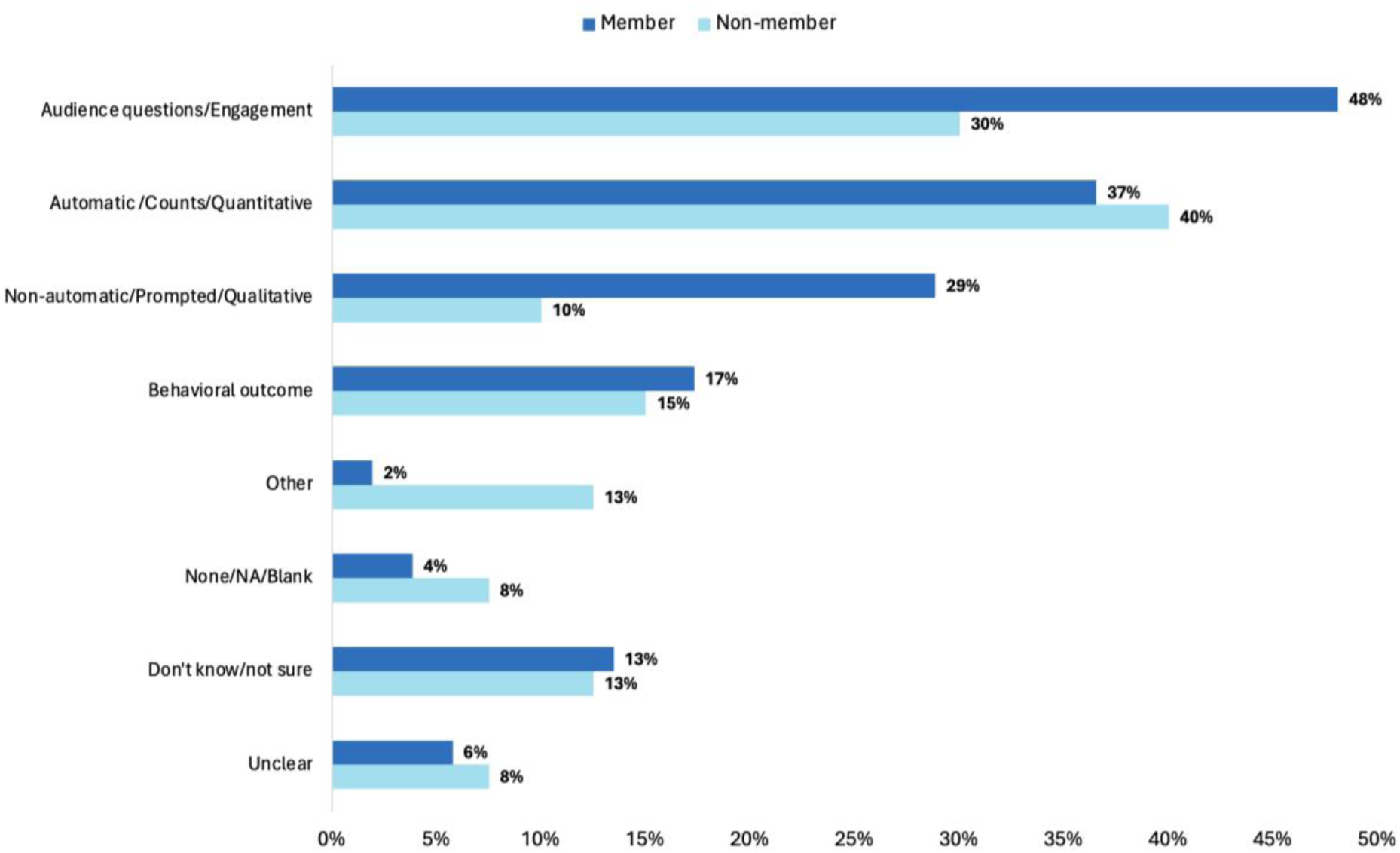
Qualitative coding of SciOSciComm-SIG member (n = 52) and non-member (*n* = 40) responses to what they felt to be the most effective metrics for measuring the success of science communication activities (bottom panel). Respondents were provided an open response text box with no character limit.

Non-members more frequently specified the intended audience for science communication efforts than members did, with 45% of non-member comments mentioning a public audience and 25% focusing a fellow specialist audience (Figure 2A). Among SIG members who specified a target audience, the majority of comments (33% of total comments) focused on the public, whereas only 2% mentioned a specialist audience (Figure 2A). Several respondents, primarily non-members, provided responses with clear meaning but failed to address the given prompt (hence their exclusion from the “Unclear” category); such responses were coded as an “Other” category.

### Science Communication Frequency Measures

When asked about the frequency of any type of public- or academic-facing science communication activities in the last 12 months, SIG member respondents indicated a significantly higher frequency of public-facing science communication activities (p = 0.008143, U = 871.5), but not academic-facing activities (p = 0.5774, U = 676.5), than non-members (Figure 3). While 29% of members and 43% of non-members indicated that they have never engaged in public-facing communication about their science (Figure 3, top panel), only 9% and 8% respectively said that about academic-facing communication (Figure 3B).

**Figure 3:**
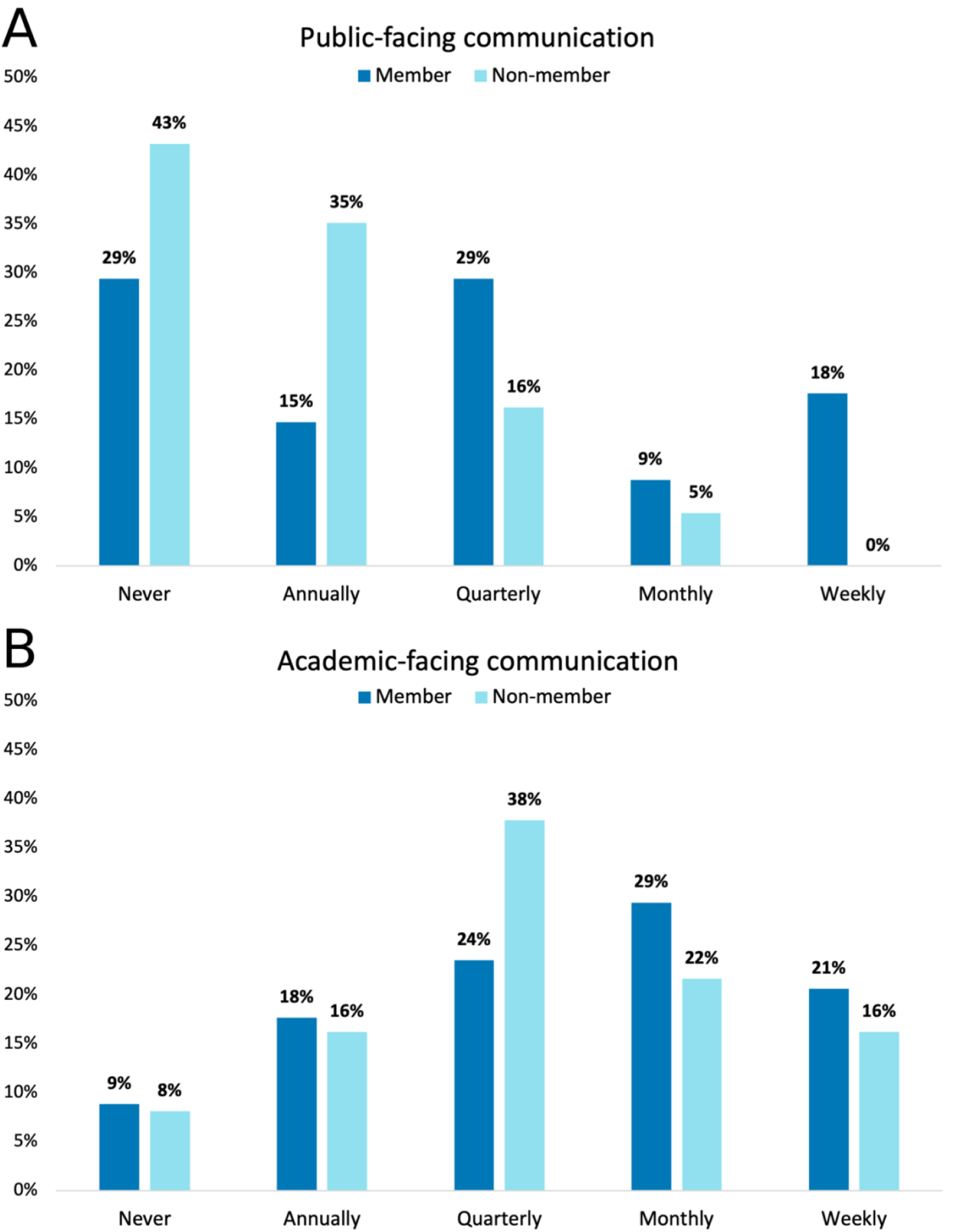
Research-affiliated member (*n* = 34) and non-member (*n* = 38) respondent frequencies of engagement in any type of public-facing (top panel) or academic-facing (bottom panel) science communication activities in the 12 months prior to survey participation.

To ascertain a more granular understanding of the types of science communication activities being performed at the NIH, we additionally asked participants about their participation in specific science communication activities that each fall under one of four subcategories. For the three public-facing science communication activity categories (Events, Traditional Media, and Online), we observed an increased frequency of “Never” responses by non-members in many of the sub-categories. For example, in the Online category, 65% of non-members indicated that their research group had never published an online video about their research in the 12 months prior to the survey, whereas only 22% of SIG members had the same response (Figure 4C). For items that had a similar prevalence of “Never” responses between groups, there was occasionally a trend toward higher frequency in the SIG member group as compared to the non-member group: referencing the “Physical media” item under the Traditional Media category, 31% of SIG members who participated in the creation of physical media related to their research at least once in the last 12 months did so on at least a monthly basis, as opposed to 6% of non-members (Figure 4B). One notable exception to the cohort participation difference seen in public-facing science communication activities is the “Festivals” item under the Events category, where non-members had more frequent participation compared to members (Figure 4A). No notable differences were identified between study groups in the academic-facing activity subcategory (Figure 4D).

**Figure 4:**
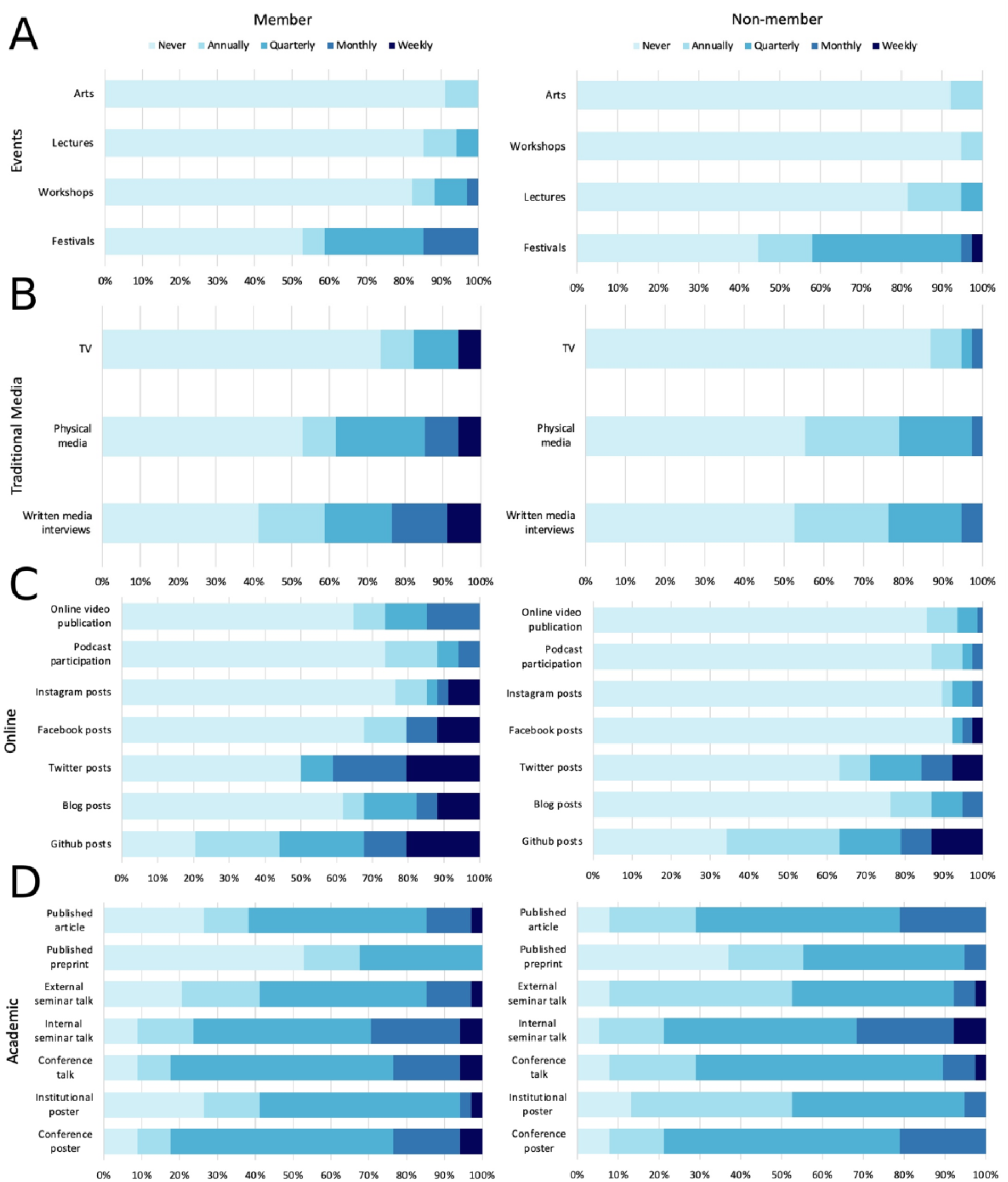
Research-affiliated member (left side panels, *n* = 34) and non-member (right side panels, *n* = 38) responses to subcategorized science communication activity frequency scales (Events [A], Traditional Media [B], Online [C], and Academic [D]) for activities occurring within the 12 months prior to survey participation.

We additionally asked participants who indicated that they are both involved in research and have engaged in at least one science communication activity in the 12 months prior to taking the survey to provide their perceived level of success of their science communication activities using a five-point Likert scale. Members and non-members alike felt that their science communication efforts were generally successful, with 79% of SIG members and 59% of non-members indicating that they perceived their science communication activities to be somewhat or very successful (Figure 5). Neutral and negative responses were generally more frequent in the non-member group than the SIG member group, together representing 21% of member and 41% of non-member responses, indicating a lower satisfaction with science communication efforts among non-members (Figure 5).

**Figure 5:**
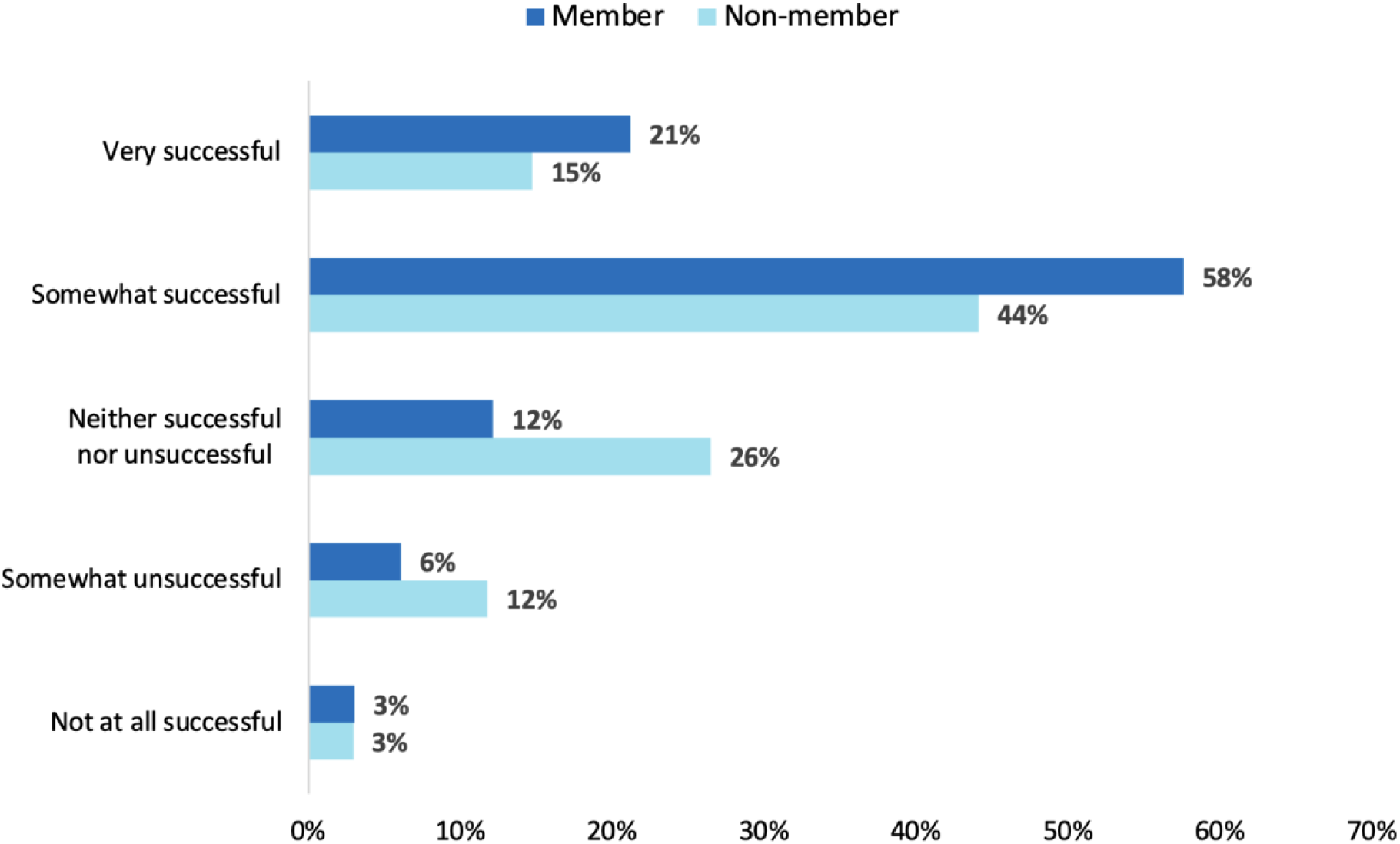
Perceived success of science communication efforts made in the past 12 months by SIG member (n = 33) and non-member (n = 34) participants.

### Barriers to Science Communication

With the understanding that researchers face a variety of well-described barriers to science communication efforts, particularly with the additional restrictions often present in government research environments, we aimed to further explore what barriers NIH researchers have faced in their science communication efforts. We approached this investigation through a mixed-methods lens, with one multiple-choice question asking about the single biggest barrier to science communication that participants have faced while working at the NIH, and another asking about experienced barriers more generally in a free-text response format. Our pre-determined options for the multiple-choice question encompassed participant responses quite well, with only 12% of members and 3% of non-members selecting the “Other” option, most of which went on to describe multiple biggest barriers or highly specific individual circumstances (Figure 6). Most participants cited administrative processes/bureaucracy or a lack of understanding/ability as the most significant experienced barrier to science communication efforts, with more non-members citing a lack of understanding/ability than members (Figure 6).

**Figure 6:**
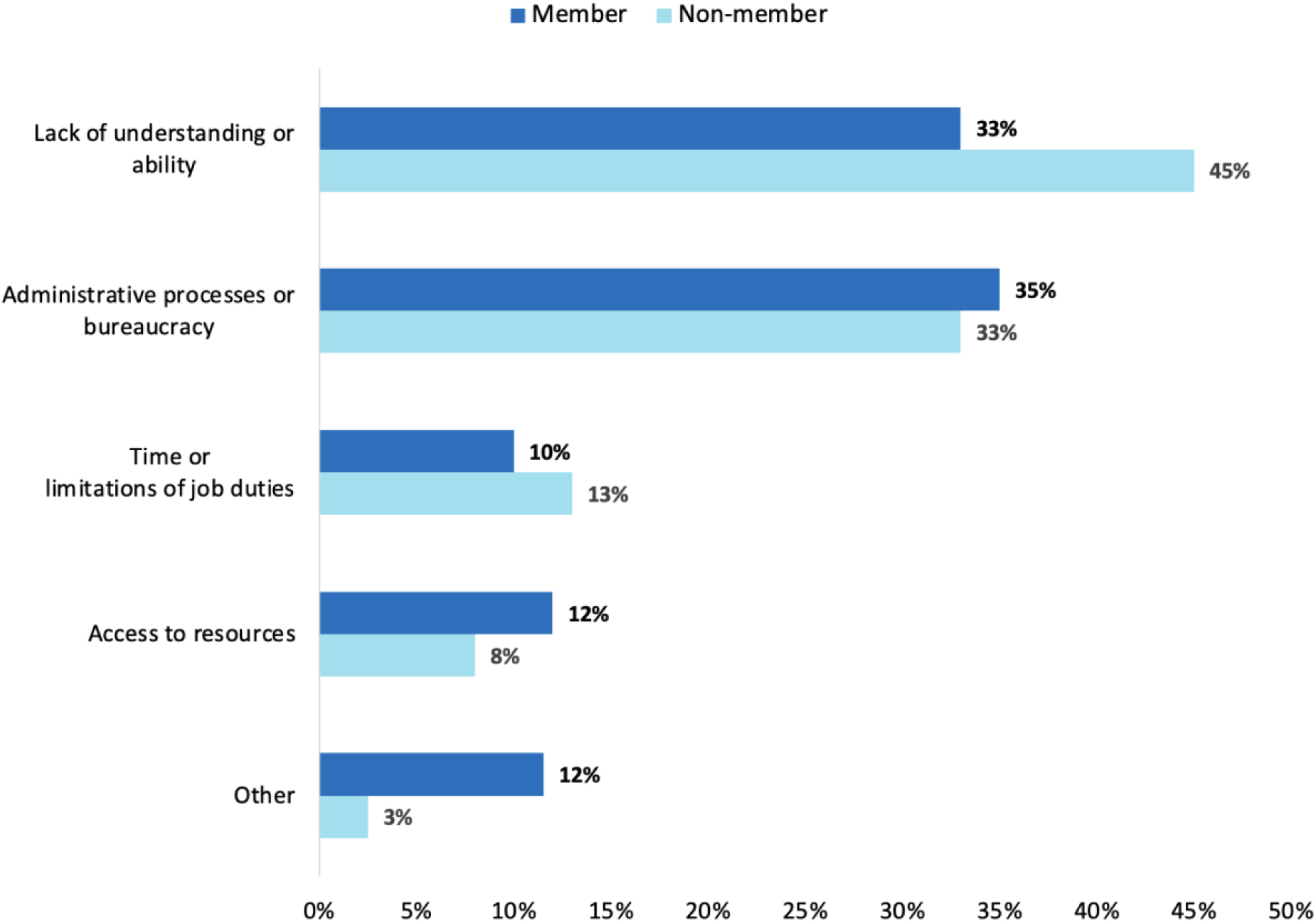
Member (*n* = 52) and non-member (n = 40) responses to a multiple-choice question asking about the biggest barrier to science communication that they have faced while employed at the NIH.

In the free-response question that asked what barriers participants have faced to science communication at the NIH in general, the responses largely reflected the options given in the multiple-choice question, although two new major themes did emerge: first, resistance or a lack of interest from colleagues in communicating science, which was often framed as a lack of responsiveness to pursuing science communication projects or a lack of enthusiasm about science communication in general. As one non-member explains:

NM05: “Lack of time, lack of guidance, lack of enthusiasm”

Similarly, one SIG member respondent expressed difficulties with engagement in science communication efforts from management or leadership:

SM46: “lack of responsiveness by [subject matter experts], lack of understanding about strategic communication in leadership”

In addition, participant SM46 also provided insight into a second novel barrier discovered in the free-response question: systemic/cultural barriers, which allude to the greater culture or conventions in academia that pose an obstacle to engaging in public-facing science communication activities across institutions and career stages. Common themes among described cultural barriers included the lack of value or focus on science communication in academia, the prioritization of writing/presenting for fellow specialists, and feelings of isolation from the general public (Table 3).

**Table 3.** Systemic/cultural barriers to science communication described by SIG member (*n* = 52) and non-member (*n* = 40) respondents in a free-text response.

| Participant ID | Comment |
| --- | --- |
| SM06 | “Too many PhDs writing for other PhDs” |
| SM24 | “Outside of publishing and to a lesser extent conferences, it's not valued” |
| NM27 | “Lack of emphasis and importance placed on science communication makes it difficult to prioritize as a trainee.” |
| NM30 | “we are isolated from the community (in the intramural environment)” |
| NM34 | “We focused on designing, executing and publishing research work, and really forget to communicate with others what we have accomplished.” |

SIG members who had attended at least one SIG meeting prior to taking the survey were additionally given the opportunity to describe what, if any, barriers to science communication that attending the SIG has helped to alleviate. Members described a range of factors related to SIG membership that helped to address barriers to science communication, including specific techniques, exposure to work being done outside of the NIH, and a feeling of community stemming from interacting with others who are experiencing similar barriers (Table 4).

**Table 4.**
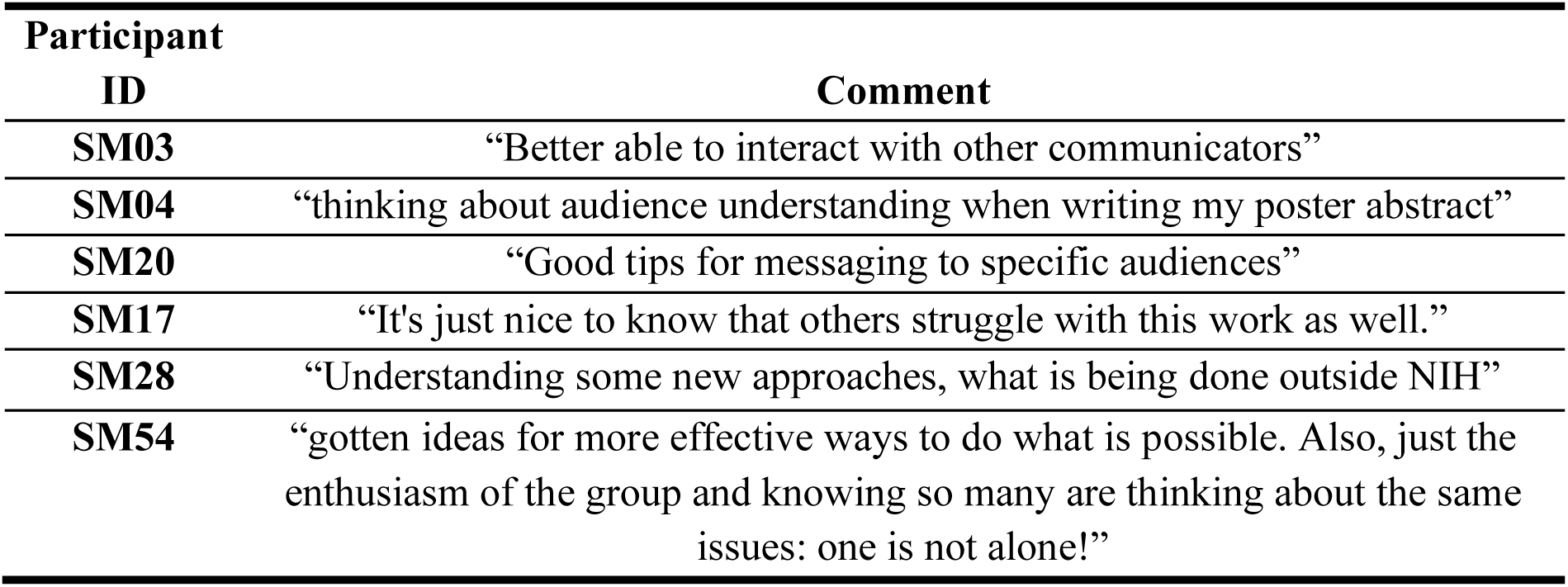
Representative free-text responses of SIG members (*n* = 52) when asked, “What, if any, barriers to science communication SciOSciComm-SIG helped to alleviate or address for you or your research group?”

### Impact of the SIG and Member Feedback

To more deeply assess the impact of the SIG on members’ work practices, we asked member participants about what, if any, techniques learned from the SIG they have applied in their own work. Members collectively described applying concepts or techniques from every one of the SIG meetings held prior to survey administration at least once, indicating a wide array of transferable skills learned from the SIG (Table 5).

**Table 5.** Representative SIG member responses (*n* = 52) to, “What knowledge, techniques, approaches, etc. highlighted in SciOSciComm-SIG meetings, if any, have you applied in your own work?”

| Participant ID | Comment |
| --- | --- |
| SM07 | “I was able to use the preprint information in my work because I work closely with publications.” |
| SM11 | “[I’ve] taken the idea of communicating uncertainty to heart and have applied it to the way [I] write papers and presentations for general audiences” |
| SM17 | “The reminder to back strong metrics and evaluation into any program.” |
| SM33 | “Making sure the facts are correct and the use of plain language” |
| SM51 | “Application of intersectionality to science communication efforts.” |

All SIG member respondents were asked to describe the impact of potential changes made to the SIG on their likelihood of future meeting attendance. SIG members seemed satisfied with the current monthly meeting frequency as well as the current proportion of general body meetings, with most responses to these items being neutral, though they were divided on the proportion of journal-club-type meetings (data not shown). Respondents were generally in favor of changes that would increase member input on SIG activities, namely diversifying speaker topics, allowing members to nominate speakers directly, and participating in polls about desired speaker topics, with all associated responses being neutral or positive (Figure 7).

**Figure 7:**
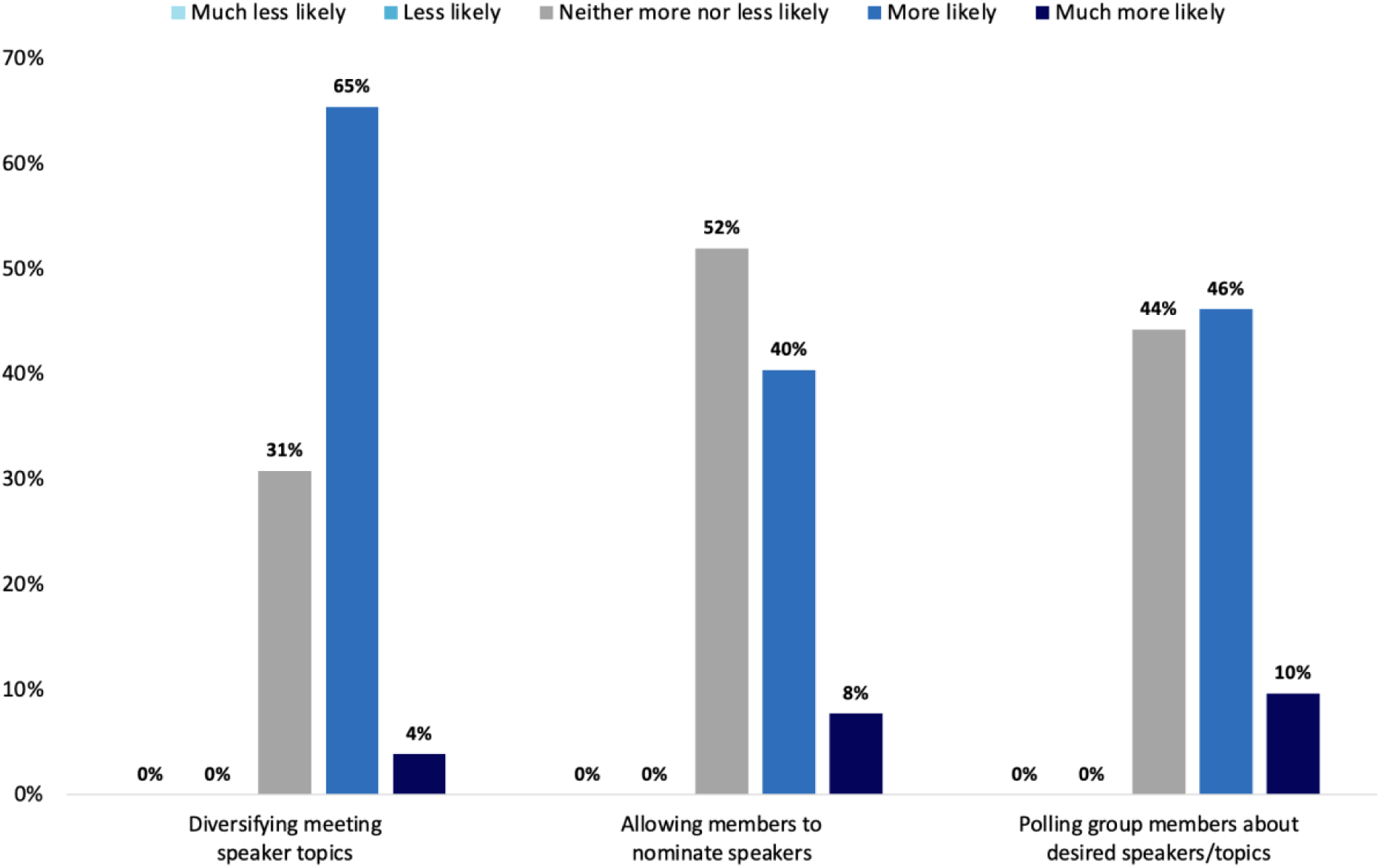
SIG member (*n* = 52) feedback on the effect of proposed changes to the SIG on personal future meeting attendance frequency.

## Discussion

Our exploratory survey found that staff and scientists at the US NIH have an interest in participating in science communication efforts, and those that have participated generally perceive their efforts to have been successful. This perceived success was present regardless of whether respondents considered themselves as members of our Science of Science Communication SIG. However, many SIG members and an even greater proportion of non-members have not participated in any type of public-facing science communication in the past year (Figure 4); therefore, their answers presumably reflect levels of success with academic audiences and methods rather than public ones. This relatively low frequency of public-facing science communication is in line with previous studies of natural scientists [Yuan, Besley, & Dudo, 2019], but poses an interesting contrast with the qualitative results in Figure 2, where non-members listed “communication with the public” as their top “most important goal of science communication.” We suggest there remains an ongoing need and appetite for both targeted training and establishment of communication opportunities for biomedical researchers, especially for multiple audiences.

We identified numerous barriers to science communication pursuits in our analysis, with the most frequent being “administrative/bureaucratic” barriers for members of our SIG, and “lack of understanding or ability” for non-members. The NIH has significantly expanded training and informational opportunities about science communication in the last decade, including “Three-Minute Talks” competitions and training provided to students at many ICOs [“Three-Minute Talks,” 2023]; the NIH Intramural Research Program’s “SciBites” videos on Instagram and “Sense About Science” podcast; regular lectures and workshops by outside and inside experts on science communication (including CG); and our own SIG as examples. Still, we found that, depending on the platform, 50->90% of our respondents indicated that they had never posted about science on social media in the past year; at the same time, 35->50% had never submitted a preprint in the same timeframe. Although working at a federal government institution presents its own challenges with bureaucracy and administrative processes, previous surveys have indicated that administrative hurdles around science communication are ubiquitous [Rose et. al, 2020]. Our respondents indicated that within the NIH, they did not feel communication was valued as part of their position (Table 3), and we felt that this fit into a greater theme of systemic and cultural barriers described by our participants. The relatively low “value” of public-facing science communication efforts and possible solutions to this problem have previously been addressed: a 2023 white paper by a coalition of scientists from US universities outlined specific steps that could be taken to recognize public-facing scholarship more clearly in promotion and tenure decisions [Ozer et. al 2023]. International groups agree; a similar theme was the first operationalizing suggestion in the Concluding Statement of the 2023 Public Communication of Science (PCST) Network Venice Symposium, which recommended specifically to “highlight the value of science communication and engagement in institutional policies, including in criteria for career advancement and promotion” [Fornetti et. al, 2023]. These changes, if implemented widely, would likely permeate to federal institutions as well, and we would therefore strongly endorse progress in this direction.

Despite their experiences of barriers, our survey found that a number of scientists at the NIH indicated monthly or even weekly posting about science on social media. In fact, members of our SIG indicated they engaged in public- and academic-facing communication weekly at roughly the same percentage frequency (Figure 3). We did not see specific reports of harassment or negative interactions during the pandemic cited as a barrier in our survey as described previously [Royan et. al, 2023], although our own experience is that changes to specific platforms and discourse have made posting on social media less frequent or enjoyable. The NIH has specified policies for private social media account use reflecting that as “a member of the NIH Community, [we] have a special responsibility to uphold the public trust” [“Guidance on Private Account Social Media Use,” 2021]. In addition, US federal employees must be aware of laws like the Hatch Act in election years [“Federal Employee Hatch Act,” 2019], which are not directly related to science communication but may still affect posts by employees.

Specifically looking at our SIG at its one-year point, we found that member satisfaction is quite high, with few suggested changes received from member respondents; the one area where members demonstrated interest was in having greater input on future speakers and meeting topics. Many members indicated that attending the SIG directly contributed to alleviating barriers to science communication, either through specific techniques or methodologies learned through attendance or through a more general mindset that was developed as part of being an active member. Similarly, members described a wide variety of skills and techniques learned from the SIG that they have gone on to apply in their own work. The SIG has seemingly gone beyond its original mission of providing a forum to learn about and discuss evidence-based science communication and has now evolved into a productive community space from which its members gain new skills, collaboratively discuss and remedy obstacles, and tangibly apply what they have learned from the group in their own work. The possibility of creating similar groups at other types of scientific institutions should be considered a promising option for a more widespread discussion on evidence-based practices in science communication.

Our study largely focused on staff who are directly involved with research, though questions on science communication and its barriers more generally as well as feedback on the SIG were open to all respondents when applicable. We did include the perspectives of communications staff and other administrative roles as they are key contributors in the relevant NIH ecosystems, setting this work apart from other studies that included only faculty or lab heads, for example. Our survey of federal government staff recapitulates themes of institutional science communication found in other types of organizations [Peters, 2022] but also shows that a group focused on the “science of science communication” can create a community of shared interest and learning in this specific environment and provide insight for those wishing to advance this interest elsewhere. Future directions for our self-evaluations include examinations of changes in knowledge, attitudes, or behavior because of the SIG and/or relevant communication activities by the NIH, which will provide further insight into the potential of the group to foster greater change in communication practices [Pellegrini, 2021].

## Acknowledgements

This research was supported by the Intramural Research Program of the National Human Genome Research Institute, National Institutes of Health.

## Authors

Please do not write anything here.

This section will be filled during the typesetting phase with the information provided in the Author Biographies field.

