## Supplementary figures and images for "Survey-Based Analysis of a Science of Science Communication Scientific Interest Group: Member Feedback and Perspectives on Science Communication"

### Supplemental Figure 1

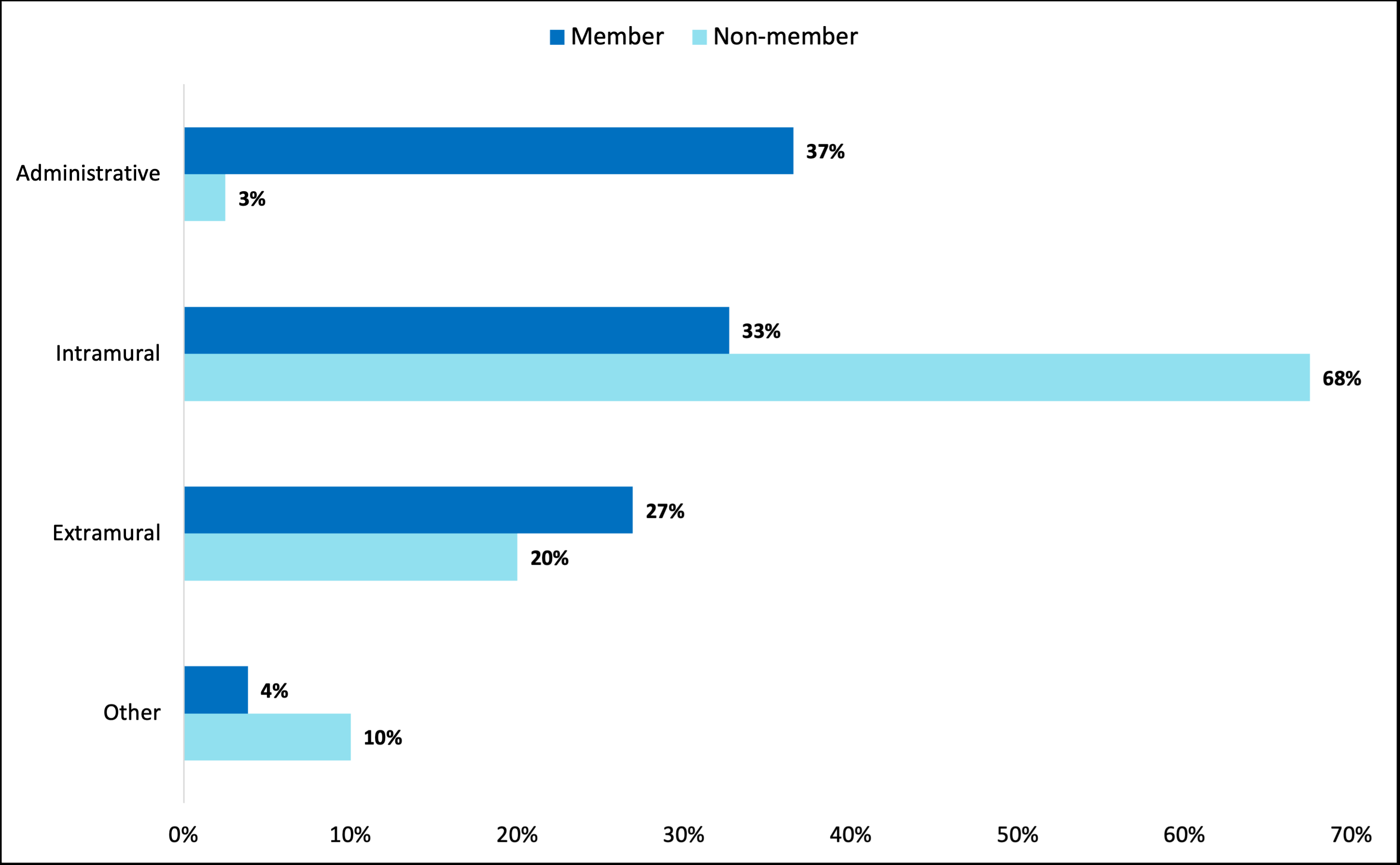

### Supplemental Figure 2

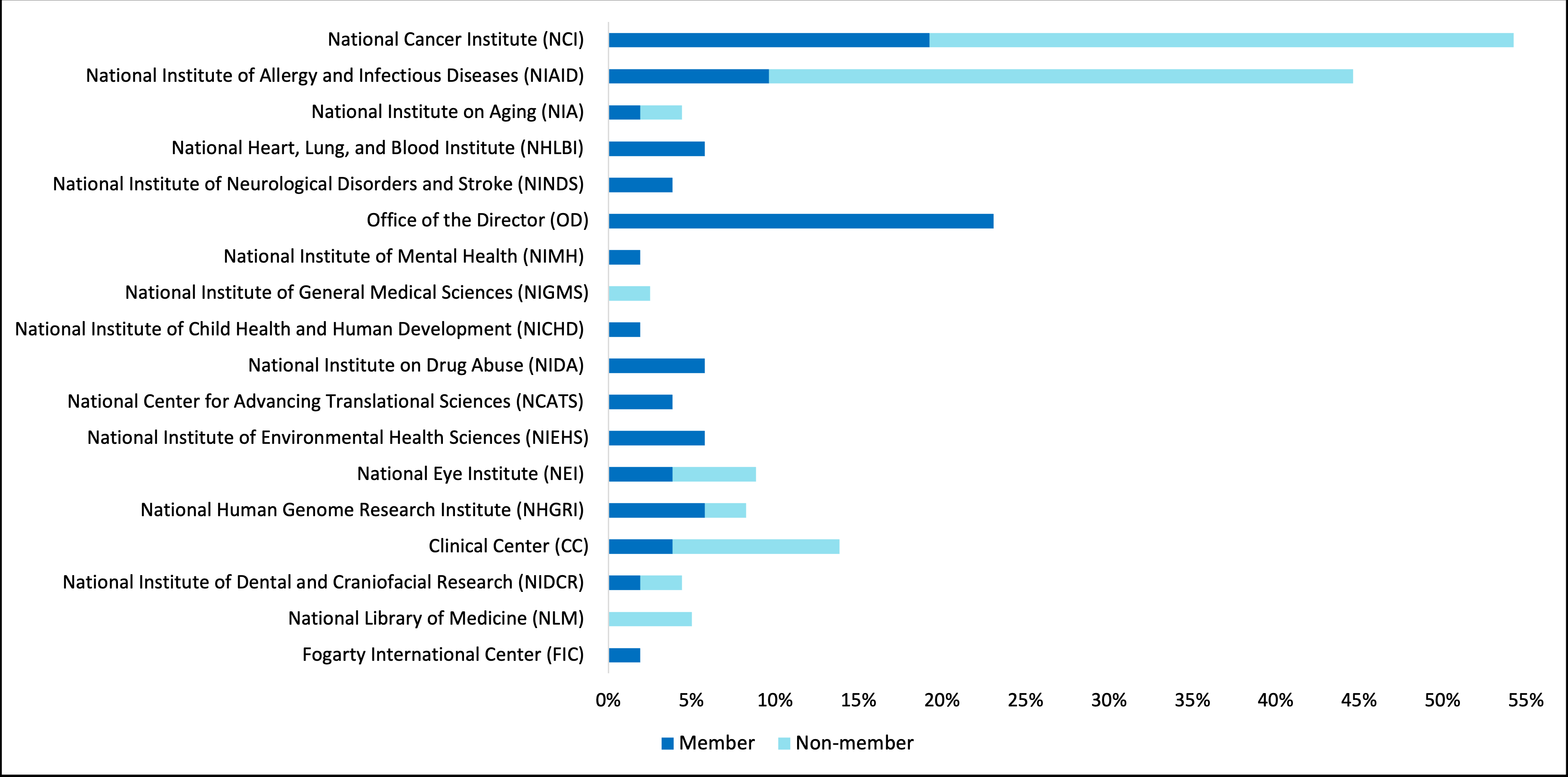
